# A highly versatile real-time quantitative RT-PCR method and sampling strategies for the accurate detection of citrus yellow vein clearing virus

**DOI:** 10.64898/2026.05.12.724569

**Authors:** María Martínez-Solsona, Ana B. Ruiz-García, Félix Morán, Beatriz Navarro, Francesco Di Serio, Melike Yurtmen, Mengji Cao, Changyong Zhou, Antonio Olmos

**Affiliations:** Instituto Valenciano de Investigaciones Agrarias (IVIA). Carretera CV-315, Km 10.7, 46113 Moncada, Valencia, Spain; Biotechnology PhD Program; Universitat Politècnica de València. Camino de Vera, s/n, 46022 Valencia, Spain; Institute for Sustainable Plant Protection, National Research Council, 70122, Bari, Italy; Biological Control Research Institute, Republic of Türkiye, Ministry of Agriculture and Forestry. Kışla Street, 01321, Yüreğir/Adana, Turkey; National Citrus Engineering Research Center, Citrus Research Institute, Southwest University, Chongqing 400712, China

**Keywords:** Citrus, Yellow vein clearing disease, CYVCV, Validated diagnostic tool, Real-time RT-qPCR, RT-dPCR, Portable real-time RT-qPCR

## Abstract

Citrus yellow vein clearing virus (CYVCV) is the causal agent of an emerging disease representing a potentially high-impact threat for citrus production. Despite remaining outside Europe for decades, CYVCV has now expanded towards two important European citrus producers, Italy and, more recently, Spain. The presence of this virus in the EPPO region represents a current threat with unpredictable and potentially devastating consequences for European citriculture. Therefore, urgent protective measures need to be taken to prevent CYVCV spread and minimize its impact. Diagnostics is a key measure in the management of viral diseases, highlighting the need for harmonized methods suitable for reliable routine detection of the currently known CYVCV diversity. In this study, an inclusive, efficient and highly sensitive real-time RT-qPCR for the detection of CYVCV in plant material and transmission vectors has been developed and validated according to EPPO standards. Moreover, the validated method has been successfully adapted to both PCR digital platforms, that allow high-sensitive absolute quantitative detection, essential in the diagnostics at low viral concentrations; and PCR portable tools, that can be applied in a real diagnostic context for on-site detection. This versatility combines standard validated performance, absolute sensitive quantitation and real on-site detection. The study has also addressed sampling strategies to support reliable molecular diagnostic performance. Our results represent an improvement in the detection of CYVCV to be applied in epidemiological studies and different real diagnostic contexts for the containment of this important citrus pathogen.

## 1. Introduction

Citrus yellow vein clearing virus (CYVCV), currently classified as *Potexvirus citriflavivenae* within the subgenus *Mandarivirus*, genus *Potexvirus*, is the causal agent of citrus yellow vein clearing disease. Since its first recognition in Pakistan, the virus has been reported from several citrus-producing regions in Asia, Europe and North America, confirming its progressive international spread (Ahlawat and Pant, 2003; Loconsole et al., 2012; Abrahamian et al., 2024). Its recent detection in Italy (Cinque et al., 2024) and Spain (Martínez-Solsona et al., 2026), has added a new phytosanitary concern for Mediterranean citriculture and highlights the need for validated and standardized diagnostic protocols for surveillance, certification and epidemiological studies.

CYVCV infects a broad range of citrus hosts, although symptom expression is highly variable. Typical symptoms on susceptible hosts include vein clearing, leaf deformation and shoot abnormalities, particularly in lemon, sour orange and lime, where disease expression is usually more evident. However, many infected plants remain asymptomatic, and symptom severity is influenced by host species, cultivar, season, temperature, viral isolate and scion–rootstock combination (Chen et al., 2014; Bin et al., 2022; Abrahamian et al., 2024; Kumar et al., 2026a). Importantly, recent experimental evidence demonstrated that CYVCV is the causal agent of spring shoot leaf curl disease in ‘Shatangju’ mandarin, indicating that its biological effects are not restricted to the classical symptomatology typically described in lemon and highlighting the need to evaluate its behaviour across different citrus hosts and production systems (Liu et al., 2025).

The epidemiological significance of CYVCV is further intensified by its host range beyond the main citrus species and by its capacity to persist in alternative hosts with little or no visible symptomatology. Natural infections have been reported in non-citrus species, including grapevine and several herbaceous hosts, which may contribute to virus maintenance in agroecosystems (Önelge et al., 2011; Afloukou and Önelge, 2020). In addition, CYVCV can be transmitted by grafting, mechanical inoculation and contaminated tools, and several insect vectors have been implicated in local spread, reinforcing concern about field dissemination once the virus is introduced into a citrus-growing area (Zhang et al., 2018a; Zhang et al., 2018b; Liu et al., 2020; Bin et al., 2026).

Accurate diagnosis is further complicated by the irregular distribution of CYVCV within infected plants. Field observations and recent experimental studies indicate that viral accumulation may differ markedly among tissues, branches and seasons, so that negative results can be obtained from infected trees when inappropriate samples are collected. In ‘Shatangju’ mandarin, for example, virus accumulation was shown to be higher in leaves during spring but to shift towards roots in winter, illustrating the importance of tissue selection and sampling time for dependable detection (Liu et al., 2025). These features make CYVCV a pathogen for which diagnostic performance depends not only on assay reliability, but also on the sampling strategy.

Over the last decade, molecular diagnostic methods have been developed. Conventional RT-PCR provided the basis for early molecular identification and remains useful for confirmatory testing and sequence-based characterization (Loconsole et al., 2012; Chen et al., 2014). Subsequent studies introduced quantitative real-time RT-qPCR (Loconsole et al., 2012; Chen et al., 2016; Liu et al., 2019; Abrahamian et al., 2024), reverse transcription droplet digital PCR (Liu et al., 2019), and isothermal approaches such as RT-LAMP (Liu et al., 2015) and RT-RPA (Gupta et al., 2024; Kumar et al., 2026b), each contributing improvements in sensitivity, quantification or operational simplicity. However, not only the increase of knowledge in viral diversity requires a continuous update of the methods, but also validation data on the diagnostic protocols, frequently absent, are essential to ensure reliable performance.

Improving CYVCV detection is not merely a technical objective but a prerequisite for effective plant health management. Validated molecular protocols according the European and Mediterranean Plant Protection Organization (EPPO) standards (PM7/98, EPPO 2025) are essential to support management measures as surveillance, sanitation programmes and to generate epidemiological data needed to understand virus spread under Mediterranean conditions, including appropriate sampling procedures in order to minimise false negatives and provide a sound basis for phytosanitary decision-making.

The present study reports the development and validation according EPPO standards of a quantitative real-time RT-qPCR assay for CYVCV detection and quantification in both plant material and transmission vectors. The protocol was designed for use not only in standard real-time qPCR platforms but also in digital PCR (dPCR) applications, aimed at absolute quantification and viral titer studies, and portable magnetic induction PCR devices to facilitate potential use under field conditions or at border inspection points. In parallel, virus distribution was investigated in infected lemon and sweet orange plants by analysing different plant tissues, in order to provide sampling strategies needed for the improvement of diagnostic reliability.

## 2. Methods

### 2.1. Plant and insect material

A total of 41 citrus trees from Spain and Italy used in this study included different cultivars of *Citrus sinensis*, *Citrus reticulata*, *Citrus clementina, Citrus aurantium, Citrus latifolia* and *Citrus limon* (listed in Supplementary Table S1). In addition, CYVCV-inoculated *Citrus volkameriana* was used as positive control and virus-free Forner Alcaide 5 (C*. reshni* Hort. Ex Tan. x *P. trifoliata*) was used as a healthy control and diluting agent. Two insect species that are common citrus vectors, *Aphis gossypii* and *Aphis spiraecola,* were collected from CYVCV-infected plants located in the Valencian Community.

### 2.2. Sample preparation and RNA purification

Plant material analyzed in this study included: leaves with petioles, young stems (without lignification), mature stems (slightly lignified), shoots (tender, including newly emerged leaves) and flowers. Plant material was collected individually and transported under refrigeration. Samples were placed in plastic bags (Bioreba, Reinach, Switzerland) and homogenized in PBS buffer at a 1:5 (w:v) ratio. Homogenization was performed manually using a hammer and resulting plant crude extracts were subsequently analyzed or kept at -80°C until further processing for RNA purification.

For plant crude extracts direct analysis, 1:100 dilution of the plant crude extract in RNase-free water (Sigma-Aldrich, Co., St. Louis, USA) was prepared. For evaluation of performance and validation assays, ten-fold dilutions were prepared using healthy plant material as a diluting agent.

For vector testing, the insects were place in microtubes in pools of 3-4 individuals and homogenized in 150 µL of PBS buffer using a micro pestle. The homogenate was centrifuged at 15,000 x g for 5 min and the supernatant was kept for further processing.

Total RNA was purified from 200 μL of plant crude extract or 100 μL of clarified insect homogenate using VANME Magnetic Universal Plant Total RNA Kit (RMA4101-C1-P3) (Vazyme Biotech Co., Jiangsu, China) in an automatic nucleic acid extraction instrument (Vazyme Biotech Co., Jiangsu, China) and following the manufacturer’s instructions. The RNA was quantified using a DeNovix DS-11 spectrophotometer (DeNovix Inc., Wilmington, DE, USA). All RNA purifications were stored at −80 °C until subsequent analysis.

### 2.3. CYVCV specific primers and probe design

For primers and probe design, 82 sequences of complete CYVCV genomes from Asia, Europe, and USA were obtained from NCBI (accessed November 2025). These sequences were aligned using MAFFT v.7 (Katoh et al., 2013) implemented in Geneious Prime 2026 software (Biomatters Ltd., Auckland, New Zealand) and manually inspected to ensure positional homology and conserved regions were studied. Two primers and a probe were designed covering a region of 176 nt: forward primer, CYVCV-PP-F1 (5′-TTAGGGGTGCCATCGAACGC -3′); reverse primer, CYVCV-PP-R2-B (5′-AGAGAAGGGGTTAGARCCRAT -3′); and probe, CYVCV-P1 (5′-FAM-TACAGCCTGACGCCGCCGACGCAC-BHQ-3′). OligoAnalyzer Tool (Integrated DNA Technologies Inc., Coralville, IA, USA) was employed for melting temperatures and secondary structure evaluation. Primers and probe were purchased from Metabion international AG (Martinsried, Bavaria, Germany).

Two additional primers were designed for cloning a region of 328 nt including the PCR target: forward primer, CYVCV-Clone-F (5’- TCATCGTCTCTGCATCAACA -3’); and CYVCV-Clone-R (5’- AGGAAGAGCATGGTGACACG –3’).

### 2.4. TaqMan Quantitative Real-Time CYVCV Detection Method

RT-qPCR assays were carried out in a QuantStudio 3 real-time PCR system (Applied Biosystems, Foster City, CA, USA), QuantStudio 5 real-time PCR system (Applied Biosystems, Foster City, CA, USA) and LightCycler 96 (Roche, Basel, Switzerland) using AgPath One-Step RT-PCR kit (Applied biosystems, Foster City, CA, USA) following the manufacturer’s instructions. The master mix contained 0.5 µM concentration of forward primer, 1.5 µM of reverse primer and 0.2 µM of the probe. The reaction mixture was carried out in a total volume of 12 µL in the case of QuantStudio 3 and QuantStudio 5 and in 20 µL in the case of LightCycler 96. All reactions contained 3 µL of RNA or diluted plant extract as template. The amplification conditions consisted of 45°C for 15 min, 95°C for 10 min, and 40 cycles of amplification (15 s at 95°C, 1 min at 60°C). The default threshold set by the machine was slightly adjusted above the noise to the linear part of the growth curve at its narrowest point, according to the manufacturer.

### 2.5. Molecular taxonomic classification of insects

For molecular taxonomic classification of insects, the *COI* gene was used, which was amplified using primers LCO1490 and HCO2198, described in PM7/129 (2) Appendix 1 (EPPO, 2021), for RNA amplification. AgPath One-Step RT-PCR kit (Applied biosystems, Foster City, CA, USA) was used. Primers were used at a concentration of 0.4 µM and 3 µL of RNA template was employed. The amplification conditions were as follows: 1 step of 45 min at 45°C; 1 step of 10 min at 95°C; 5 steps of 30 s at 94°C, 30 s at 45°C and 1 min; 35 steps of 30 s at 94°C, 1 min at 51°C and 1 min at 60°C; with a final step of 10 min at 60°C. The amplicons were Sanger sequenced and species-assigned by BLASTn comparison.

### 2.6. Validation of the Real-Time qPCR Method

The RT-qPCR method developed in this study was evaluated following EPPO standards, listed in PM 7/98, Appendix 5, table A7 (EPPO, 2025).

#### 2.6.1. Evaluation of the Sensitivity

For absolute quantitation, a standard curve was constructed with CYVCV RNA *in vitro* transcribed from plasmid DNA bearing a viral fragment containing the PCR target sequence. Briefly, viral sequence was amplified from RNA purified from naturally infected CYVCV sample using primers CYVCV-Clone-F and CYVCV-Clone-R (see section 2.3) and AgPath One-Step RT-PCR kit (Applied biosystems, Foster City, CA, USA) following the manufacturer’s instructions. The reaction mixture was composed of 0.5 µM of each primer and was carried out in a total volume of 25 µL containing 2 µL of template RNA using an Applied Biosystems™ Veriti™ Thermal Cycler (Fisher Scientific, Loughborough, Leicestershire, UK). The amplification conditions consisted of a reverse transcription step at 45°C for 45 min and a denaturation step of 95°C for 10 min, followed by 35 cycles of amplification (15 s at 95°C and 1 min 30 s at 60°C) and a final step of 10 min at 60°C. The 328 bp PCR product was purified using the mi-PCR Purification Kit (metabion international AG, Martinsried, Germany), subsequently ligated into the pGEM®-T Easy Vector (Promega Corporation, Madison, WI, USA) according to the manufacturer’s protocol and transformed into *Escherichia coli* NZY5α (NZYtech, Lisboa, Portugal) competent cells. Positive transformants were selected on LB agar plates supplemented with ampicillin (50 µg/mL), IPTG (0.134 mM) and X-Gal (0.5 mg/mL). Plates were incubated at 37°C for 24 hours. Plasmid DNA was extracted using the mi-Plasmid Miniprep Kit (metabion international AG, Martinsried, Germany) according to the manufacturer’s instructions and quantified with a DeNovix DS-11 spectrophotometer (DeNovix Inc., Wilmington, DE, USA). Subsequently, the plasmid was linearized with *SalI* restriction enzyme and 1200 ng of linearized DNA used as a template for *in vitro* transcription with T7 RNA polymerase (Takara Bio Inc., Kusatsu, Japan), following the manufacturer’s recommendations and adding 1µL of Ribolock (Thermo Fisher Scientific, Waltham, MA, USA). The reaction was incubated at 42 °C for 2 hours. Residual DNA was then removed by digestion with RQ1 RNase-free DNase (Promega Corporation, Madison, WI, USA) at 37 °C for 30 min. Finally, the synthesized RNA was purified using the spin columns provided in the Plant/Fungi Total RNA Purification Kit (Norgen Biotek Corporation, Thorold, ON, Canada) and quantified.

The number of RNA transcripts was calculated using the formula: ssRNA = (µg of ssRNA × 10^6^)/(320.47× Nb), where 320.47 corresponds to the average molecular weight of a ribonucleotide and Nb represents the total number of bases in the transcript. The transcript copy number was subsequently estimated using Avogadro’s constant (6.023 × 10^23^ molecules/mol) (Avogadro, 1811).

Three replicates of serial dilutions from 1.2 × 10^8^ to 1.2 RNA copies/µL were prepared and used to generate the standard curve. The amplification efficiency of the calibration curve was calculated by the slope, according to the mathematical formula: amplification efficiency = [10 ^(−1/slope)^] −1 (Bustin, 2000).

The analytical sensitivity in plant material was evaluated using either crude extracts or purified RNA obtained from naturally CYVCV-infected sample (LBC1). Ten-fold serial dilutions of plant crude extracts in healthy crude extracts were analyzed in triplicates. The number of viral targets was determined by interpolation in the standard curve.

#### 2.6.2. Evaluation of the Specificity

The inclusivity of the developed method was first evaluated *in silico* using 213 CYVCV sequences retrieved from NCBI (accessed March 2026) together with 24 nearly complete genomes from Spanish isolates sequenced by our group. The dataset comprised isolates from China (n = 80), India (n = 28), Spain (n = 27), Turkey (n = 21), the USA (n = 21), Iran (n = 14), Italy (n = 12), South Korea (n = 8), and Pakistan (n = 2). Sequences were aligned using the Geneious Prime 2026 software (Biomatters Ltd., Auckland, New Zealand), and mismatches in the binding regions of CYVCV primers and probes previously reported in the literature were evaluated for critical mismatching, according to Morán et al. (2023). Inclusivity was also confirmed experimentally with 32 CYVCV-infected samples from different growing regions from Spain and 1 CYVCV-infected sample from Italy. Exclusivity was assessed by testing 8 CYVCV-free citrus samples infected by other viruses and viroids commonly present in this crop as determined by HTS.

#### 2.6.3. Evaluation of selectivity

For selectivity, different citrus species (*Citrus limon*, *Citrus volkameriana*, *Citrus reticulata*, *Citrus clementina, Citrus sinensis, Citrus aurantium* and *Citrus latifolia*) and cultivars (Clemenvilla, Clemenules, Navelate, Navelina, Valencia Late, Fino and Verna), listed in Supplementary Table S1. In addition, different plant tissues were used, including flowers, leaves with petioles, shoots, young stems, and mature stems. The method was also tested on common citrus vectors such as *Aphis gossypii* and *Aphis spiraecola*.

#### 2.6.4. Evaluation of the Repeatability and Reproducibility

For the evaluation of repeatability and reproducibility, 6 naturally CYVCV-infected samples with a relatively low viral concentration were selected and analyzed using purified RNA (samples 248.1, 248.2 and LBC2) or crude extracts (samples 303.3, 295.1 and LBC1). Samples were analyzed in three independent experiments carried out on different days, by different operators and three different thermal cyclers (see section 2.4). Three replicates, giving a total of nine final experimental replicates for each sample, were tested.

### 2.7. RT-dPCR assay

RT-dPCR was carried out in a PCR digital Applied Biosystems QuantStudio Absolute Q system (Applied Biosystems, Thermo Fisher Scientific, Waltham, MA, USA) using Absolute Q™ 1-Step RT-dPCR Master Mix kit (Applied Biosystems, Thermo Fisher Scientific, Waltham, MA, USA), following the manufacturer’s instructions. The same primers and probe were used using the concentrations mentioned above (sections 2.3, 2.4). All reactions contained 2.7 µL of RNA template in a reaction volume of 9 µL. Amplification conditions consisted of 10 min for reverse transcription at 55°C, a denaturing step at 96°C for 10 min, and 40 cycles of amplification at 96°C for 5 s and 60°C for 30 s.

For the plate configuration, each sample and its ten-fold dilutions were considered a group, and a negative control included. The threshold was determined automatically. Once the total concentration of copies/µL in the sample was determined, the total number of copies was calculated based on that result.

### 2.8. Magnetic induction RT-qPCR system

For magnetic induction RT-qPCR, Q (Quantabio, Beverly, MA, USA) system was employed with qScript XLT One-Step RT-qPCR ToughMix (Quantabio, Beverly, MA, USA) following the manufacturer’s instructions.

The same primers and probe were used at the concentrations mentioned above (sections 2.3, 2.4). The reaction was carried out in a volume of 20 µL, using 3 µL of the RNA template or diluted plant extract. The amplification conditions were as follows: 45°C for 10 min, 95°C for 1 min, and 30 cycles of amplification (3 s at 95°C, 10 s at 60°C). The default threshold set by the machine was slightly adjusted above the noise to the linear part of the growth curve at its narrowest point, according to the manufacturer.

### 2.9. Evaluation of sampling influence on CYVCV detection

For the first approach, two lemon trees naturally infected by CYVCV (LBC1 and LBC2) were selected, and two symptomatic and two asymptomatic branches, were taken from each tree. Leaves and petioles were separated from the stems, and both tissues processed and analyzed separately in triplicates, yielding a total of 6 samples per tissue type analyzed.

For the second approach, three lemon trees and three sweet orange trees (LBC3, LBC4, LBC5, OBC1, OBC2, OBC3) naturally infected by CYVCV were selected. From each tree, 7-8 asymptomatic branches were taken and divided into 5 tissue types: flowers, leaves with petioles, young stems, mature stems and shoots, which were processed and analyzed separately (35–40 samples analyzed per tree).

### 2.10. Statistical analysis

Contingency tables and Pearson’s chi-square test were used to analyze the association between CYVCV detection and categorical variables. Detection was classified as positive or negative; tissue type included five categories (flowers, leaves with petioles, shoots, young stems and mature stems), and host species included two categories (lemon and sweet orange). Separate analyses were performed for tissue type and host species. Statistical analyses were performed using JASP version 0.96 (JASP Team, 2026).

## 3. Results

### 3.1. Development of a real-time RT-qPCR method for CYVCV detection

In order to design a highly inclusive CYVCV detection method, all CYVCV full genomes available in the databases, including the recently identified Spanish isolates (Martínez-Solsona et al., 2026), were used. A total of 82 genomes were aligned, and the most conserved regions studied. This analysis led to the RdRp gene, in which a 176 nt PCR-target region was selected. Amplification of this genomic fragment was achieved *in silico* in all genomes by two primers: primer CYVCV-PP-F1 and primer CYVCV-PP-R2B. In addition, a single TaqMan probe showing specificity to all CYVCV sequences was found, CYVCV-P1. The performance of these primers and probe towards CYVCV detection was tested *in vitro* for 33 CYVCV infected samples from different geographic origins: 32 samples from different growing regions in Spain and 1 sample from Italy (listed in Supplementary Table S1). All CYVCV infected samples were successfully detected by the real-time assay developed.

### 3.2. Validation of the diagnostic method according to EPPO standards

#### 3.2.1. Sensitivity

Analytical sensitivity was evaluated by absolute quantitation using *in vitro* transcribed CYVCV RNA. Three replicates of ten-fold serial dilutions of the quantified target RNA containing from 3.6 x 10^8^ up to 3.6 molecules were tested. The method was able to detect CYVCV in all the serial dilutions tested, thus showing a limit of detection of around 3 viral copies. However, a standard curve with a high amplification efficiency of 99.56% (coefficient of correlation, R^2^=0.997) was achieved for a quantitation range between 30 and 3 x 10^7^ molecules (Figure 1A).

**Figure 1.**
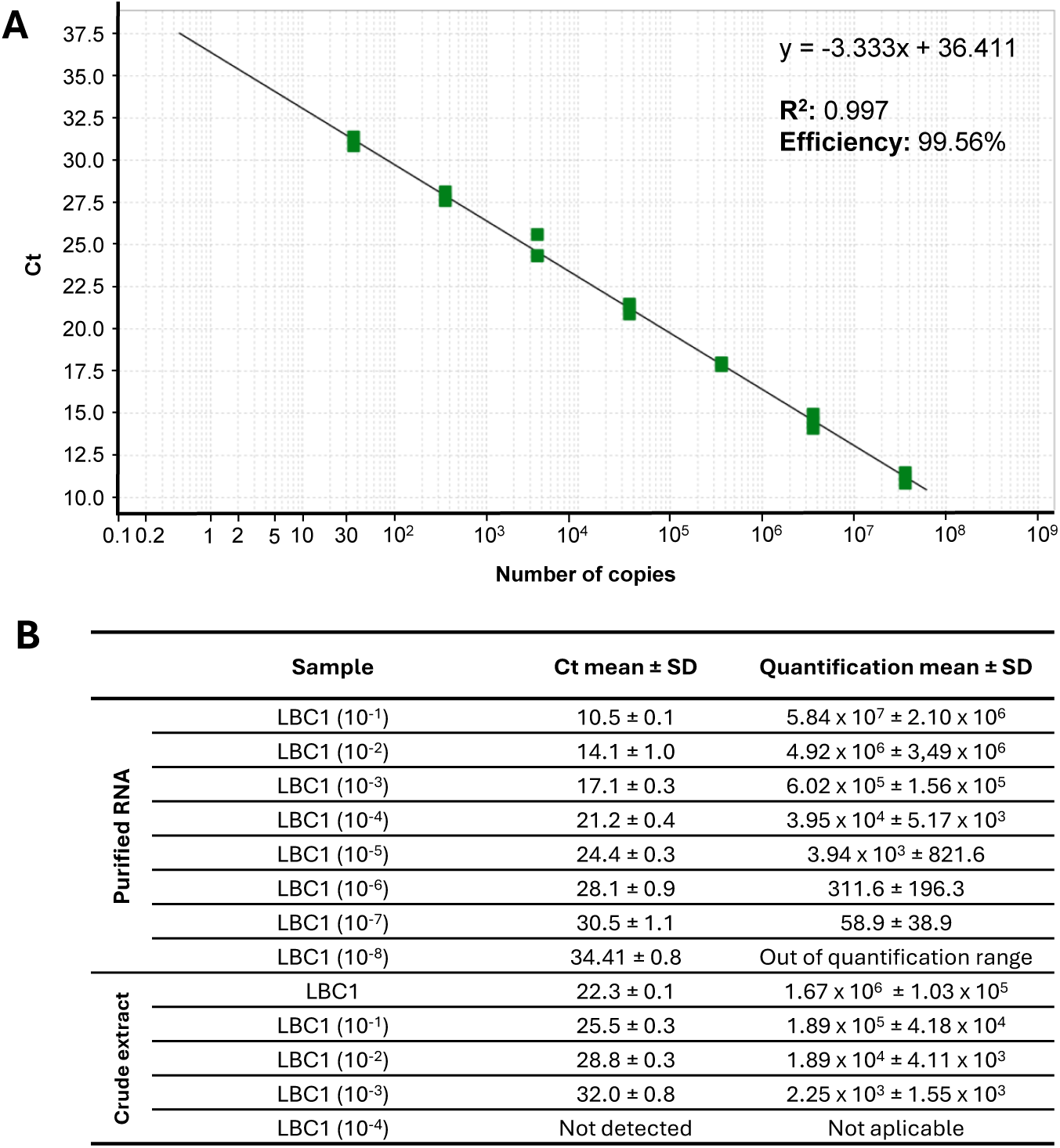
Analytical sensitivity of the RT-qPCR method developed in this study for the detection of CYVCV. **(A)** Standard curve for the absolute quantitation of CYVCV. The Ct values of three replicates of CYVCV RNA transcripts ten-fold serial dilutions are plotted along with the number of copies. Mathematical equation for the standard quantitation curve, correlation coefficient (R²) and efficiency are indicated. **(B)** Absolute quantitation of ten-fold serial dilutions of purified RNA and crude extract of CYVCV-infected sample (LBC1). Mean Ct values and SD, as well as mean total number of copies quantified and SD are indicated.

In parallel, three replicates of ten-fold serial dilutions of both crude extract and purified RNA from one CYVCV citrus infected sample (LBC1) were analyzed and quantified. According to the standard curve, the detection limit for this naturally infected plant material was established at 59 viral copies for purified RNA and 2,250 viral copies for plant crude extract. (Figure 1B).

Taken together, these results show that the real-time RT-qPCR method developed in this study is able to detect CYVCV with a very high sensitivity, allowing an accurate quantification range of the viral titer from 30 to 3 x 10^7^ RNA copies, with a limit of detection around 3 viral RNA molecules. The sensitivity for viral RNA isolated from infected plant material was established at a higher limit of detection of around 60 copies. Interestingly, even using plant crude extract, the method is able to detect around 2,200 viral copies, avoiding a nucleic acid purification step.

#### 3.2.2. Analytical specificity and selectivity

Analytical specificity of the technique was evaluated in terms of inclusivity and exclusivity. Inclusivity was first evaluated *in silico* during the primers/probe designing step, that perfectly aligned with all the CYVCV full genomes available to date, with a matching percentage at each nucleotide position of more than 99%. In addition, during the validation step a more in depth analysis showed an inclusive *in silico* performance on 213 CYVCV full and partial sequences available in the databases, including sequences from China, India, Spain, Turkey, the USA, Iran, Italy, South Korea and Pakistan, as well as 24 nearly complete Spanish genomes recently sequenced by our group (unpublished results).

Inclusivity was also tested experimentally with CYVCV infected samples from different geographic origins, as mentioned above, including several growing regions from Spain and one from Italy. All samples tested positive for CYVCV, demonstrating the inclusive detection ability of the technique.

Exclusivity was evaluated by the analysis of 8 CYVCV-free citrus samples naturally infected by other citrus common viruses/viroids, which virome had been previously determined by HTS (Table 1). All samples tested negative for CYVCV, indicating the absence of any unspecific reaction and thus showing the high exclusivity of the technique.

**Table 1.** CYVCV-free samples used for evaluating exclusivity of the RT-qPCR method. The selected samples were analyzed by HTS in a previous study (Martinez-Solsona et al., 2026) and are infected by several common viruses and viroids in citrus, listed in this table.

| Sample ID | Virus/Viroid Detected |
| --- | --- |
| 256.5 | CTV genotype VT, CTV genotype T30 |
| 260.5 | CTV genotype RB, CVEV, CDVd, CBLVd, HSVd, CEVd |
| 264.5 | CVEV, CTV genotype T30 |
| 267.6 | CiVA, CPsV, CTV genotype RB, CTV genotype VT, CTV genotype T30, CEVd, HSVd, CBLVd |
| 283.6 | CTV genotype T3, CTV genotype RB, CTV genotype VT |
| 314.3 | CTV genotype RB, CTV genotype VT |
| 314.4 | CTV genotype VT, CTV genotype T30 |
| 315.2 | CTV genotype RB, CTV genotype VT |

CTV: Citrus tristeza virus; CVEV: Citrus vein enation virus; CDVd: Citrus dwarfing viroid; CBLVd: Citrus bent leaf viroid; HSVd: Hop stunt viroid; CEVd: Citrus exocortis viroid; CiVA: Citrus virus A; CPsV: Citrus psorosis virus.

The selectivity of the technique was also evaluated by analyzing different plant matrices, including different citrus species: lemon, sour orange, sweet orange, lime and mandarin; different tissues of lemon and sweet orange trees, shoots, stems, leaves and flowers; as well as two different species of aphids, *Aphis gossypii* and *Aphis spiraecola*. Despite the differences in viral distribution among different citrus species/tissues, our method successfully detected CYVCV in all the matrices tested.

#### 3.2.3. Repeatability and reproducibility

Finally, to assess the performance of the technique in diagnostic laboratories, repeatability and reproducibility of the results were evaluated. Six CYVCV infected samples were tested using either plant crude extracts (three samples) or purified RNA (three samples) in three independent experiments carried out by different operators, using different instruments and testing on different days. All samples analyzed in each independent experiment were tested in triplicates, yielding nine technical repeats per sample.

Full diagnostic agreement between all replicates and repetitions was obtained, as the virus was detected in all the replicates of the infected samples, independently of the operator, the instrument used and the day of analysis (Table 2). The maximum coefficient of variation among independent assays (different operator, equipment and day) was 2.16% for plant crude extract and 5.18% for RNA purified samples. Diagnostic results were consistent in both RNA samples and plant crude extracts, showing a high level of repeatability and reproducibility.

**Table 2.** Repeatability and reproducibility of the RT-qPCR method. Six samples were analyzed in a total of nine technical repeats in three independent experiments carried out by three different operators using three different thermal cyclers (three repeats per instrument/operator), testing both crude extracts (three samples) and purified RNA (three samples). For every sample, mean Ct and SD were calculated for different instruments used in this study. In addition, the coefficient of variation (CV) between independent experiments is shown for each sample.

|  | Sample ID | Quant Studio 3 | Quant Studio 5 | Roche | CV (%) |
| --- | --- | --- | --- | --- | --- |
| Plant extract | 303.3 | 25.59 ± 0.20 | 25.57 ± 0.55 | 26.17 ± 0.65 | 2.07 |
|  | LBC1 | 25.35 ± 0.13 | 25.12 ± 0.21 | 26.18 ± 0.31 | 2.05 |
|  | 291.5 | 25.70 ± 0.16 | 26.86 ± 1.01 | 26.86 ± 0.18 | 2.96 |
| Purified RNA | 248.1 | 19.98 ± 0.22 | 19.73 ± 0.07 | 21.91 ± 0.08 | 5.18 |
|  | 248.2 | 19.41 ± 0.32 | 19.22 ± 0.12 | 21.28 ± 0.23 | 5.08 |
|  | LBC2 | 28.94 ± 0.11 | 29.63 ± 0.33 | 31.33 ± 0.56 | 3.73 |

### 3.3. Application of the method to digital PCR platforms: absolute quantitation

The real-time RT-qPCR detection method designed and validated in this study was applied for detection of CYVCV by dPCR, allowing high-sensitive detection and absolute quantitation.

Two asymptomatic mandarin samples naturally infected by CYVCV with a relatively low viral titter were used for this analysis. Four 10-fold serial dilutions from 10^-2^ to 10^-5^ were tested by RT-dPCR. The results of this experiment are shown in Figure 2. Efficient detection of CYVCV was achieved in both samples by RT-dPCR using the primers and probe designed and validated in this study (Figure 2A, 2B). In addition, absolute quantitation of CYVCV established a limit of detection between 2.07 (sample 248.1, dilution 10^-5^) and 35.64 (sample 248.2, dilution 10^-5^) viral copies. These samples were tested in parallel by real-time RT-qPCR. A positive detection was obtained for all the serial dilutions quantified by RT-dPCR except sample 248.1, dilution 10^-5^ (Figure 2A, 2B), thus confirming the high analytical sensitivity of the real-time RT-PCR evaluated by a standard curve.

**Figure 2.**
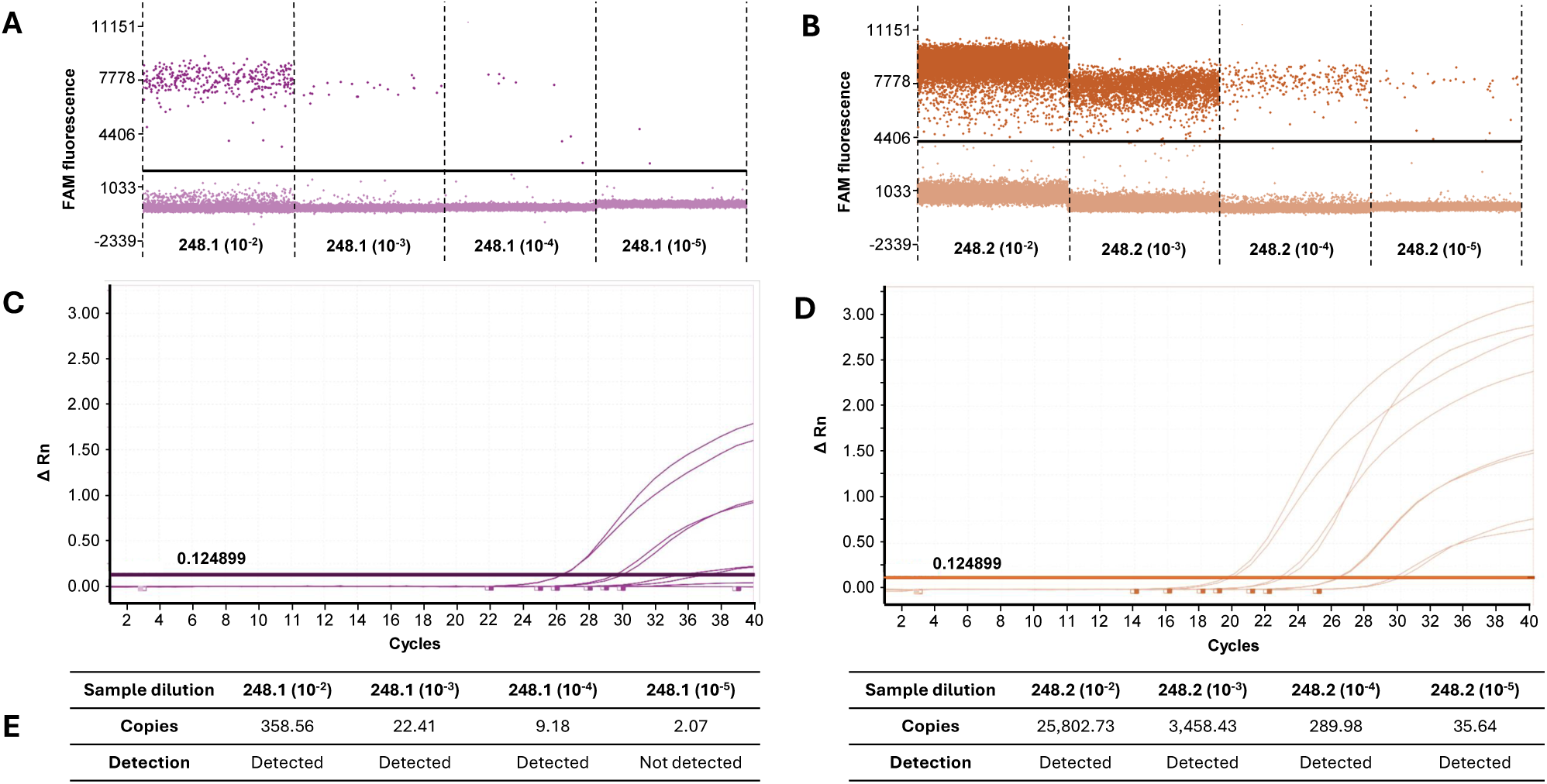
Performance of the RT-PCR assay in a digital PCR platform. Fluorescence amplitude plot corresponding to RT-dPCR analysis of ten-fold serial dilutions of sample 248.1 **(A)** and 248.2 **(B)**. Each point represents an individual partition after amplification. Positive partitions were identified by fluorescence signals above the established threshold (black line). **(C-D)** RT-qPCR analysis of the same samples. **(E)** Absolute CYVCV quantitation in total number of copies detected of all samples analyzed. The results of standard RT-qPCR analysis are shown, indicating detection or non-detection of the virus.

### 3.4. Application of the method to portable PCR platforms: on-site detection

Our CYVCV detection method was also applied in a magnetically induced PCR platform which technology allows the performance in a portable compact equipment susceptible to being used for on-site detection in border post, nursery or field conditions. To this end, the six citrus samples previously tested for repeatability and reproducibility, were tested for CYVCV detection in both purified RNA (three samples) and plant crude extracts (three samples) by magnetically induced RT-qPCR. All samples were analyzed in triplicate. In order to achieve a fast performance, suitable for an on-site detection protocol, a shorter RT-qPCR program including only 30 instead of 40 cycles of amplification was applied.

The results of this experiment are shown in Figure 3. Five out of the six samples gave a positive detection result, with similar Ct values to those obtained by standard real-time RT-qPCR reaction. Only one of the six samples (sample LBC2) resulted in a false negative result, due to a low viral titter. It is important to note that the mean Ct value for this sample when analyzed by standard real-time RT-qPCR was 29.95 ± 1.23 (Table 2) and therefore out of the amplification assay set up in the magnetic induced PCR platform.

**Figure 3.**
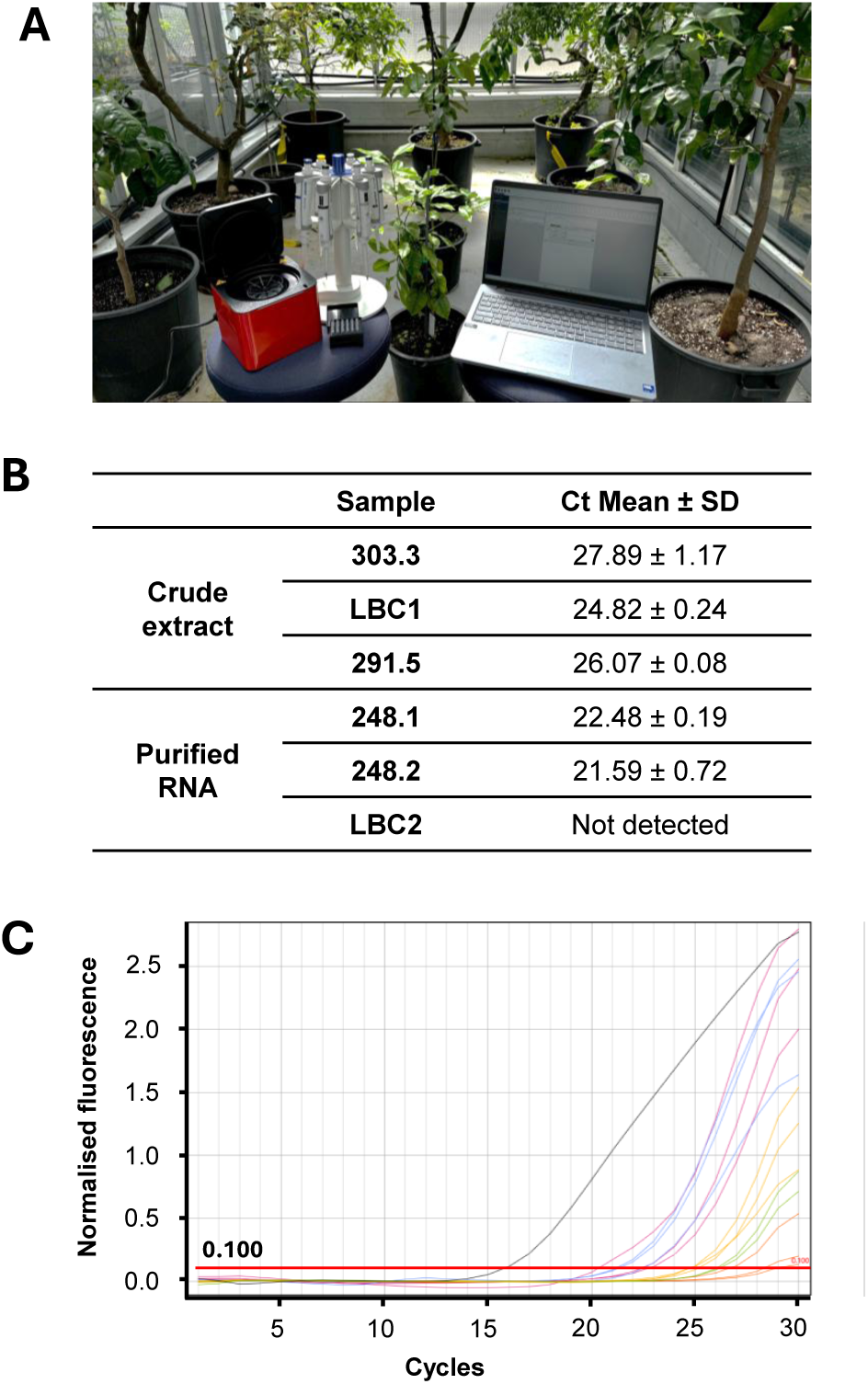
Performance of the RT-qPCR assay in a magnetic induction PCR portable platform. Six CYVCV infected samples were analyzed in a portable magnetic induction PCR device **(A)**, testing purified RNA (three samples) and crude extracts (three samples). A shorter amplification reaction of 49 minutes, including 30 instead of 40 amplification cycles was used. Mean Ct values and SD **(B)** and amplification plots **(C)** are shown.

These results demonstrate the ability of the developed method to be applied in on-site diagnostics conditions, using portable compact platforms.

### 3.5. Sampling strategies for improved CYVCV detection by real-time RT-qPCR

To evaluate the effect of sampling on CYVCV detection we first compared symptomatic and asymptomatic plant material of two CYVCV infected lemon trees showing typical symptoms in many, but not all, of their branches. Both stems and leaves with petioles were collected and analyzed in triplicate from two symptomatic and two asymptomatic branches from each plant (Figure 4A).

**Figure 4.**
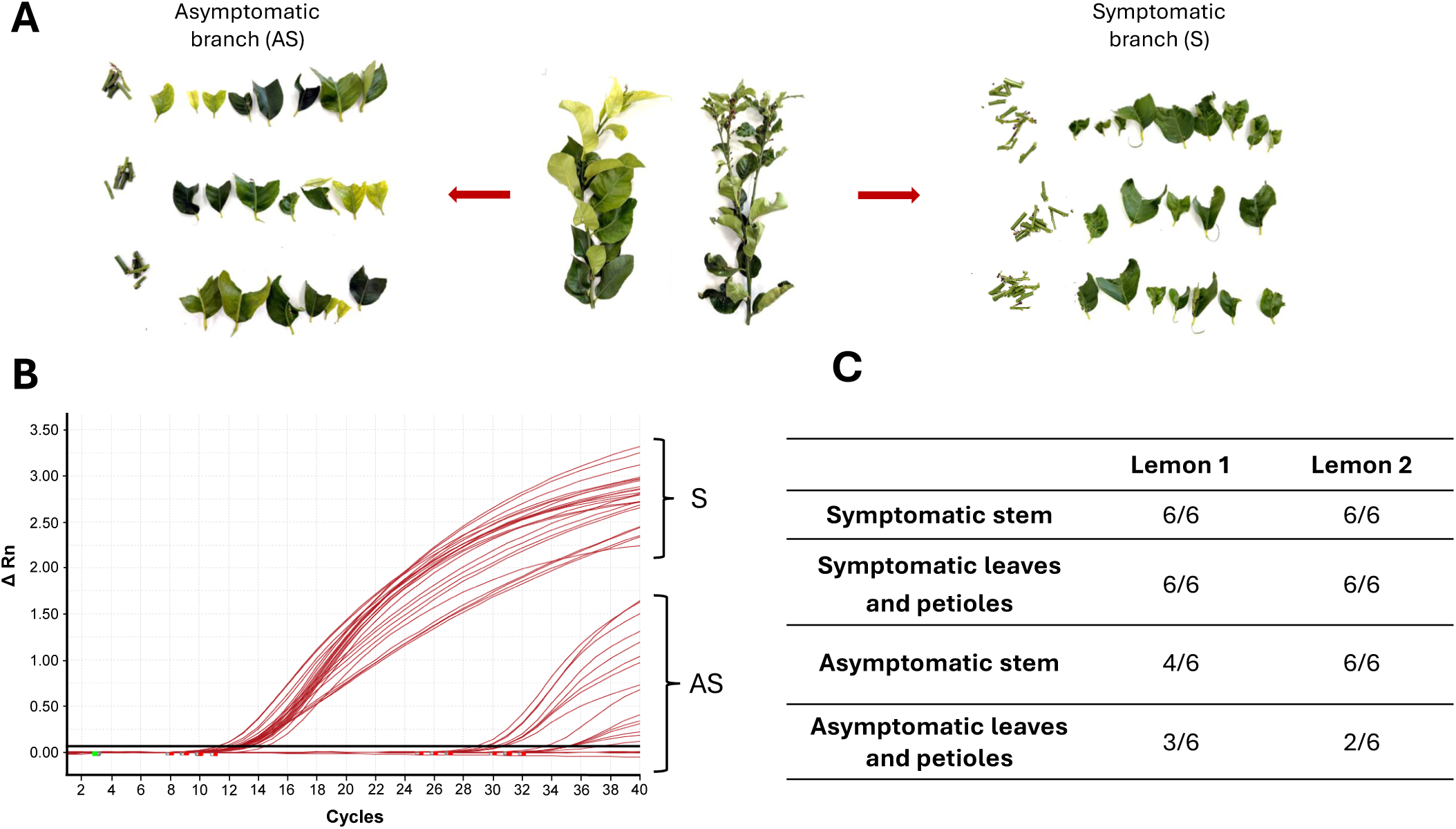
CYVCV detection efficiency comparison between symptomatic and asymptomatic branches of CYVCV infected lemon trees. **(A)** Symptomatic (S) and asymptomatic (AS) branches collected from the same CYVCV infected lemon trees were divided into stems and leaves with petioles. **(B)** Real-time RT-qPCR analysis of three replicates of the stems and leaves sampled. Amplification curves belonging to symptomatic (S) or asymptomatic (AS) branches are indicated. **(C)** CYVCV detection rate.

Although CYVCV was detected in both symptomatic and asymptomatic branches, the results of this experiment showed critical differences in the amplification signal obtained for these two types of material (Figure 4B). CYVCV detection was much less efficient in asymptomatic material, probably due to a lower viral titer. In addition, in asymptomatic material the detection was more efficient in stem tissue than in leaf tissue. These differences were not found between stems and leaves of symptomatic branches (Figure 4C).

Taking into account that CYVCV infection often remains asymptomatic in several citrus species, and with the aim of determining the best tissue to be tested for a CYVCV efficient detection, a comparison was made between different plant material collected from lemon and sweet orange trees located in the same orchard, and therefore in the same climatic and agronomic conditions. Three symptomatic lemon trees and three asymptomatic sweet orange trees were selected for this purpose, all of them infected by CYVCV.

Samples from 7-8 asymptomatic branches around the tree were collected from different tissues: shoots, young stems, mature stems, leaves with petioles and flowers (Figure 5A). Thus, 35-40 asymptomatic samples of each of the six trees (7-8 samples of each of the 5 different tissues per tree) were analyzed for CYVCV detection (Figure 5B).

**Figure 5.**
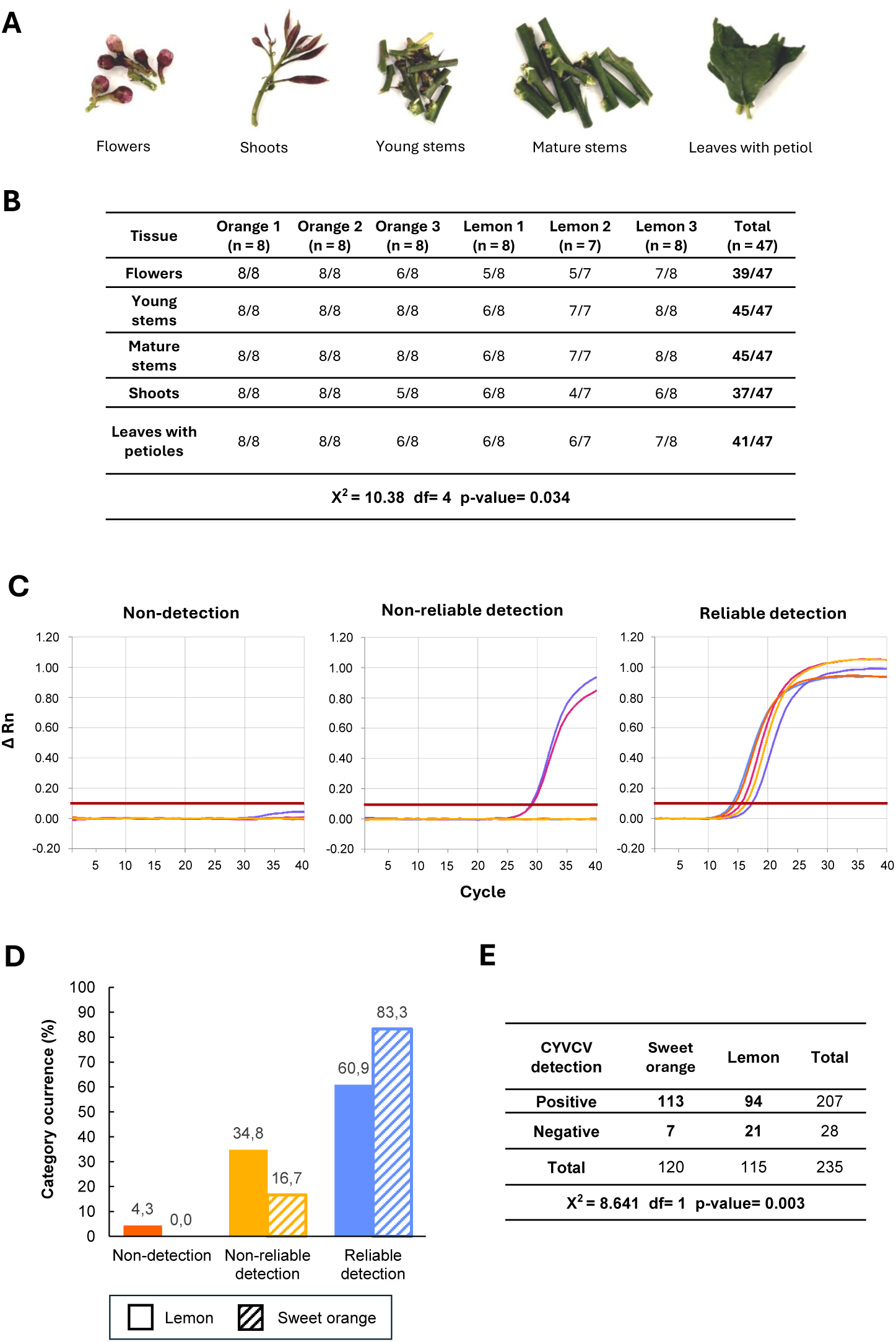
Sampling effect on CYVCV detection in sweet orange and lemon trees. **(A)** Five different tissues sampled from asymptomatic branches of sweet orange and lemon trees infected by CYVCV: flowers, shoots, young stems, mature stems and leaves with petioles. A total of 7-8 asymptomatic branches were selected from 3 sweet orange and 3 lemon trees from the same orchard. All 5 types of different tissues were sampled from each branch, yielding 35-40 samples per tree (7-8 samples for each of the 5 tissues) that were analyzed individually. **(B)** CYVCV detection rate for each tree and tissue. Chi-squared, df and p-value for the statistic comparison between total detection rate per tissue is indicated. **(C)** Qualitative diagnostic categories used to classify the overall detection efficiency achieved for every branch: non-detection (CYVCV not detected in any of the tissues); non-reliable detection (CYVCV detected only in some of the tissues); and reliable detection (CYVCV detected in all the tissues). **(D)** Percentage of diagnostic categories occurrence in sweet orange and lemon. **(E)** Contingency table for CYVCV detection rate in sweet orange and lemon. Chi-squared, df and p-value for the statistic comparison between total detection rate in both citrus species.

Overall, intriguing differences in the detection efficiency across samples were observed, as some samples resulted in a high CYVCV detection signal whereas others gave a weaker positive or even a negative testing signal. For a qualitative assessment of this sample heterogenicity, three diagnostic categories were considered in order to classify branches sampled in terms of the overall detection efficiency achieved: i) “*non-detection”*, for those branches with a negative result for all the tissues tested; ii) “*non-reliable detection”*, for those branches that gave a positive signal, sometimes close to the limit of detection, for some but not all the tested tissues; and *“reliable detection”*, for those branches that resulted in a strong positive detection independently of the tissue analyzed (Figure 5C). As a result of this classification, differences were observed between lemon and orange trees. Most of the orange branches tested lied under the *“reliable detection”* category (20/24, 83.33 %), with only occasional occurrence of *“non-reliable detection”* (4/24, 16.67 %) category. A different situation was found in lemon trees, in which low efficiency detection categories, *“non-reliable detection”* (8/23, 34.78 %) or even *“non-detection”* (1/23, 4.35 %) were more frequent (Figure 5D). These results suggest the occurrence of a more uneven viral distribution in lemon than in orange trees.

A quantitative analysis of CYVCV detection efficiency comparing different tissues was also performed. The proportion of samples testing positive for CYVCV was calculated for each of the tissues evaluated in both lemon and orange trees (Figure 5B). Young and mature stems were the tissues with a higher efficiency of CYVCV detection (24/24 and 21/23, 100 % and 91.30 %, in sweet orange and lemon, respectively), in contrast to flowers, shoots and leaves with petioles, which gave a higher rate of false negative results (Figure 5B). These differences in the testing performance of different tissues were evaluated statistically as significant by chi-squared test. In addition, significant differences statistically supported were found when sweet orange and lemon were compared (Figure 5E) supporting a more uneven viral distribution in this citrus species.

## 4. Discussion

CYVCV has recently become a phytosanitary concern of relevance for European citriculture, and as such, it is included in the EPPO Alert List and under pest risk assessment evaluation. Moreover, its recent detections in Italy and Spain, indicate that the virus can no longer be considered an exotic pest but a real threat with the potential for establishment and spread in the Mediterranean basin (Cinque et al., 2024; Martínez-Solsona et al., 2026). The current situation in Europe is particularly concerning because insect vectors are already present in the EPPO region and therefore natural spread can be expected. Under these circumstances, urgent protection measures are required for the control and containment of the virus. In this sense, reliable diagnostic methods are essential for sanitary selection, certification, surveillance and epidemiology. Therefore, harmonized methods validated according to EPPO diagnostic standards (EPPO, 2025), including analytical sensitivity, analytical specificity, selectivity, repeatability and reproducibility, are necessary.

CYVCV phylogenetic analyses have shown the genetic diversity of this virus identifying two major groups: the East Asia group, further subdivided into eight subgroups, and the South Asia, Middle East and North America group (Sun and Yokomi, 2024). More recent European findings expand this genetic diversity, with the Spanish isolates grouping with the latter lineage, and a complete isolate from Lazio (Italy) not clustering clearly within either of the two major groups. Overall, CYVCV genetic diversity has implications in the detection of this pathogen and must be considered for the development of highly inclusive diagnostic methods.

Several CYVCV detection methods have been reported in the literature. Among them, real-time RT-qPCR techniques (Loconsole et al., 2012; Chen et al., 2016; Liu et al., 2019; Abrahamian et al., 2024) are the best candidates to perform as gold standard diagnostic tools. However, a preliminar *in silico* evaluation of such methods in the context of the new CYVCV genetic knowledge, especially the recently identified Spanish genomes, showed the need of updating their inclusivity, as critical mismatches in primers and probe that could somehow limit virus detection were found. More importantly, previous studies reporting real-time CYVCV detection techniques do not provide validation data, which is a requirement for harmonization and standardization of diagnostic protocols.

On the other hand, several studies have shown that CYVCV infection is often symptomless in many citrus hosts (Zhou et al., 2017; Abrahamia et al., 2024), In addition, previous studies have reported an irregular distribution of the virus among plant tissues and variations associated with the season or phenological stage (Liu et al., 2020; Liu et al., 2025; Kumar et al., 2026a). These biological features have an important effect on the viral load present in a citrus sample and thus can be critical for reliable detection. Therefore, both a high sensitivity detection method as well as an appropriate sampling strategy are particularly relevant for accurate and efficient CYVCV diagnosis.

Taking these requirements into account, the main goal of this study was the development of a robust diagnostic protocol for the management and control of CYVCV. This protocol includes not only a validated highly inclusive and sensitive real-time RT-qPCR method but also sampling guidance for improved detection. In addition, the method has been designed for a broaden applicability to digital and portable platforms, extending its use for the detection of CYVCV under specific diagnostic contexts, such as very low viral titer, particularly relevant in propagation material and transmission vectors; or rapid on-site testing that might be required in nurseries, field or border inspection.

The real-time RT-qPCR CYVCV diagnostic method developed in this study targets a highly conserved region within the RdRp gene, covering all the genetic diversity known to date, including all CYVCV genomic sequences available in public databases, together with 24 Spanish isolates sequenced by our group (unpublished results). The inclusivity of the method has been evaluated both *in silico*, demonstrating its potential ability to detect all currently known CYVCV genetic diversity worldwide; and experimentally using CYVCV-infected samples representing the genetic diversity currently present in the EPPO region, demonstrating its ability to detect isolates from different geographic locations of Spain and Italy. Thus, this method represents an important contribution to the availability of highly inclusive CYVCV diagnostic tools. These results, together with the absence of unspecific cross-reactions with other common citrus viruses and viroids, demonstrate the high analytical specificity of our method.

We have also demonstrated the high analytical sensitivity of the method developed in this study. The assay was able to detect 3 RNA transcript copies by the real-time RT-qPCR, with a reproducible quantification limit of approximately 30 viral copies, being 60 copies the limit of detection in plant material. The sensitivity was even higher when the method was adapted to a digital platform, allowing the detection of 2 copies of the viral genome. These values make the method suitable for the detection of low viral titers allowing the analysis of samples in which the amount of target RNA may be limited, including propagation material and insect vectors.

The method has been further validated for performance on different matrices, showing an efficient CYVCV detection on samples from different citrus species, sweet and sour orange, mandarin, lime and lemon; as well as different citrus tissues, stems, leaves, shoots and flowers; and insect vectors, *Aphis gossypii* and *Aphis spiraecola*. This high selectivity is also relevant to support broad citrus protection measures and epidemiological studies.

Finally, the high repeatability and reproducibility shown by the method applied in replicates by different operators and using different instruments highlight the robustness of the technique for harmonization and transferability.

During the final stage of validation of the CYVCV diagnostic method developed in this study, another RT-qPCR assay was published (Osman et al., 2026). This technique shows *in silico* a high inclusivity and has been validated according to Minimum Information for Publication of Quantitative Real-Time PCR Experiments (MIQE) guidelines. This study also represents a useful contribution to the current need for updated and validated CYVCV diagnostic tools.

As mentioned above, and in agreement with previous observations, CYVCV is usually not evenly distributed within infected trees, which can represent an important diagnostic limitation. In the present study, this aspect has been further analyzed under Mediterranean conditions by comparing diagnostic performance when testing different plant materials collected from two different citrus species, sweet orange, generally bearing asymptomatic CYVCV infections, and lemon, frequently showing typical disease symptoms. The analysis of a total of 235 samples of flowers, young stems, mature stems, shoots and leaves with petioles from 6 CYVCV infected trees (3 sweet orange and 3 lemon trees) cultivated in the same agronomic and climatic conditions showed statistically significant differences in the diagnosis efficiency of the virus between both citrus species, and also between different tissues. First of all, our results show that the diagnostic efficiency is higher in sweet orange, where consistency in the results obtained from different tissues (94.16%) is higher than in lemon (81.74%), showing this citrus species a high percentage of false negative results. In addition, our results indicate that diagnostic performance is highly influenced by the tissue analyzed, being stems a more suitable plant material to be tested for efficient CYVCV detection. This sampling effect should be considered and incorporated into official diagnostic protocols for which, based on our results we recommend testing young or mature stems from 4-8 branches from the tree, in order to minimize false negative results occurrence.

The versatility of our CYVCV real-time RT-qPCR assay towards digital and magnetically induced portable PCR platforms adds also practical value required in specific real diagnostic contexts. Digital platforms are particularly useful as they allow absolute quantitation and provide reliable detection for low-titer samples and therefore can be applied for the analysis of propagation material in sanitation and certification programs; as well as in epidemiological studies, for the detection of the virus in transmission vectors. In this sense, it is worth highlighting the high analytical sensitivity of the method developed that has allowed the detection of 2 target copies. This high sensitivity, very close to the limit of detection, and therefore compromised in real-time standard platforms, can however be confidently achieved in the digital application. On the other hand, portable platforms may facilitate rapid on-site testing in nurseries, field inspections or border control contexts. Our method has also shown an efficient performance in this application. It is important to note that because these platforms are intended for rapid *in situ* testing a shorter amplification reaction for the detection on plant crude extract has been applied, resulting in a lower limit of detection of approximately 2,000 viral copies, which is nevertheless at the optimum range for a rapid detection applied in preliminary screenings.

## 5. Conclusion

A new real-time RT-qPCR diagnostic method for CYVCV detection was developed using updated sequence information, including newly identified Spanish isolates, and validated according to EPPO standards. The assay showed excellent analytical performance and applicability across different matrices and platforms, supporting its use as part of a harmonized diagnostic framework for CYVCV in the EPPO region. In addition, the study demonstrates that diagnostic reliability relies on sampling strategies as well. Stem tissue resulted in the most suitable matrices for efficient CYVCV detection, which should be sampled instead of shoots, flowers or leaf tissue to improve diagnostic performance. The uneven distribution of the virus in the tree, especially in lemon, reveals that negative results from single asymptomatic branches should be interpreted with caution. Consequently, in view of this irregular distribution and taking the high sensitivity of the method into account, the results support the use of pooled stem material collected from four to eight branches distributed around the canopy. Overall, this work combines a reliable validated assay with practical sampling recommendations to provide a complete diagnostic protocol to support the management and control of CYVCV.

---

**Supplementary Table S1.** Citrus trees used in this study.

| Sample ID | Host | Cultivar | Rootstock | Province | Location | Collection date | CYVCV detection |
| --- | --- | --- | --- | --- | --- | --- | --- |
| 246.2 | *Citrus sinensis* | Navelina | Carrizo | Valencia | Genovés | September 2025 | Detected |
| 246.5 | *Citrus sinensis* | Navelina | Carrizo | Valencia | Genovés | September 2025 | Detected |
| 247.1 | *Citrus sinensis* | Navelina | Carrizo | Valencia | Rótova | September 2025 | Detected |
| 247.2 | *Citrus sinensis* | Navelina | Carrizo | Valencia | Rótova | September 2025 | Detected |
| 247.3 | *Citrus sinensis* | Navelina | Carrizo | Valencia | Rótova | September 2025 | Detected |
| 247.5 | *Citrus sinensis* | Navelina | Carrizo | Valencia | Rótova | September 2025 | Detected |
| 247.6 | *Citrus sinensis* | Navelina | Carrizo | Valencia | Rótova | September 2025 | Detected |
| 248.1 | *Citrus reticulata* | Clemenvilla | Carrizo | Valencia | Gandía | September 2025 | Detected |
| 248.2 | *Citrus reticulata* | Clemenvilla | Carrizo | Valencia | Gandía | September 2025 | Detected |
| 248.5 | *Citrus reticulata* | Clemenvilla | Carrizo | Valencia | Gandía | September 2025 | Detected |
| 248.6 | *Citrus reticulata* | Clemenvilla | Carrizo | Valencia | Gandía | September 2025 | Detected |
| 249.1 | *Citrus clementina* | Clemenules | Carrizo | Castellón | Sant Jordi | September 2025 | Detected |
| 256.5 | *Citrus clementina* | Clemenules | FA5 | Valencia | Sagunto | October 2025 | No detected |
| 260.5 | *Citrus sinensis* | Washington Navel | Troyer | Valencia | Chiva | October 2025 | No detected |
| 264.6 | *Citrus sinensis* | Navelina | Macrophylla | Alicante | Formentera del Segura | October 2025 | No detected |
| 267.6 | *Citrus clementina* | Cultifort | Citrumelo | Castellón | Onda | October 2025 | No detected |
| 278.1 | *Citrus sinensis* | Navelina | Carrizo | Valencia | Carcaixent | October 2025 | Detected |
| 278.3 | *Citrus sinensis* | Navelina | Carrizo | Valencia | Carcaixent | October 2025 | Detected |
| 278.4 | *Citrus sinensis* | Navelina | Carrizo | Valencia | Carcaixent | October 2025 | Detected |
| 278.5 | *Citrus sinensis* | Navelina | Carrizo | Valencia | Carcaixent | October 2025 | Detected |
| 283.6 | *Citrus sinensis* | Navelina | Carrizo | Valencia | Pobla Llarga | October 2025 | No detected |
| 291.5 | *Citrus clementina* | Clemenules | Carrizo | Castellón | Betxí | October 2025 | Detected |
| 298.1 | *Citrus sinensis* | Navelate | Carrizo | Castellón | Vinaroz | October 2025 | Detected |
| 303.1 | *Citrus sinensis* | Valencia Late | Carrizo | Valencia | Benifairó de la Valldigna | September 2025 | Detected |
| 303.3 | *Citrus sinensis* | Valencia Late | Carrizo | Valencia | Benifairó de la Valldigna | September 2025 | Detected |
| 303.5 | *Citrus sinensis* | Valencia Late | Carrizo | Valencia | Benifairó de la Valldigna | September 2025 | Detected |
| 303.6 | *Citrus sinensis* | Valencia Late | Carrizo | Valencia | Benifairó de la Valldigna | September 2025 | Detected |
| 314.3 | *Citrus clementina* | Neufina | Carrizo | Valencia | Carcaixent | November 2025 | No detected |
| 314.4 | *Citrus clementina* | Nero | Carrizo | Valencia | Carcaixent | November 2025 | No detected |
| 315.2 | *Citrus clementina* | Neufina | FA5 | Valencia | Moncada | November 2025 | No detected |
| Italy lemon | *Citrus limon* (L.)<br>Burm.f. | Unknown | Unknown | Campania | Palma<br>Campania | May 2024 | Detected |
| LBC1 | *Citrus limon* | Verna | Unknown | Castellón | Benicarló | February 2026 | Detected |
| LBC2 | *Citrus limon* | Verna | Unknown | Castellón | Benicarló | February 2026 | Detected |
| LBC3 | *Citrus limon* | Verna | Unknown | Castellón | Benicarló | March 2026 | Detected |
| LBC4 | *Citrus limon* | Verna | Unknown | Castellón | Benicarló | March 2026 | Detected |
| LBC5 | *Citrus limon* | Verna | Unknown | Castellón | Benicarló | March 2026 | Detected |
| OBC1 | *Citrus sinensis* | Unknown | Unknown | Castellón | Benicarló | March 2026 | Detected |
| OBC2 | *Citrus sinensis* | Unknown | Unknown | Castellón | Benicarló | March 2026 | Detected |
| OBC3 | *Citrus sinensis* | Unknown | Unknown | Castellón | Benicarló | March 2026 | Detected |
| 335 | *Citrus aurantium* | Unknown | Unknown | Castellón | Sant Jordi | December 2025 | Detected |
| 344.4 | *Citrus latifolia* | Unknown | Unknown | Castellón | Vinaroz | February 2026 | Detected |

